# Morphological and molecular diversity of rice blast isolates from Terai foothills of the Himalayas

**DOI:** 10.64898/2026.07.18.739344

**Authors:** Susmita Jha, Yeluru Mohan Babu, Kumar Sathiyaseelan, Jh Ashwini, Suparna Das, Rosa Sanchez-Lucas

## Abstract

Rice blast disease, caused by *Magnaporthe oryzae* (formerly as *Pyricularia grisea*), is a major threat to rice production globally, causing devastating yield losses up to 30-50% of rice production annually. Here, we analysed 48 isolates collected from rice fields in the Terai foothills of the Himalayas, India, to assess pathogen occurrence in a new agroecological zone, potentially influenced by climate change and favourable environmental conditions. All isolates were screened for virulence and pathogenicity, and two highly virulent isolates (UBKV1 and UBKV2) were selected for detailed characterization of their morphological, growth, and genetic variability. Significant differences (p-value=4.16 × 10⁻⁵) were observed in conidial dimensions, with UBKV2 producing larger spores (33.61 µm) compared to UBKV1 (28.07 µm). In terms of growth, mycelial biomass (fresh weight) and sporulation intensity was also higher in UBKV2 (22.98 g and 2315.5) than UBKV1 (15.82 g, and 1812.3) when they grew under the same conditions. Distinct colony growth patterns were observed on different media, particularly on Mathur’s medium and Rice Straw Extract Dextrose Oatmeal Agar, where UBKV2 exhibited suppressed growth and unique pigmentation. Phylogenetic analysis of the ITS region revealed sequence similarities ranging from 95.11% to 100% among the isolates. UBKV2 showed closer genetic relatedness to isolates from Odisha (96.95–97.23%) than to UBKV1 (95.57%), highlighting significant genetic differentiation. These findings demonstrate substantial morphological, cultural, and genetic variation within *M. oryzae* populations in the Terai foothills, providing important insights into pathogen evolution, virulence mechanisms, and implications for region-specific resistance breeding strategies.

## 1. Introduction

Rice (*Oryza sativa*) is one of the world’s most vital crops, feeding over half of the global population (Birla et al., 2017) particularly across Asia. It is also a major source of income for millions of smallholder farmers, making its sustainable production essential for both food security and rural livelihoods. With global population growth, rice demand is expected to rise sharply, requiring substantial production increases by 2050 (FAO, 2022a, b). This highlights the need for innovative farming, particularly disease management, to sustain yields. In Asiatic and South American countries, rice is integral to culture and agriculture (Bellwood, 2023, Bandumula et al., 2018). The Terai foothills of the Himalayas form an important rice-growing agroecological zone that shares climatic and agricultural characteristics with eastern Nepal and adjoining rice-producing regions of South Asia. This interconnected landscape supports extensive rice cultivation but may also facilitate the emergence and spread of rice blast populations across regional production systems (Durai et al., 2015; Das et al., 2020). Thus, maintaining both productivity and biodiversity of locally adapted rice varieties is therefore essential for regional and global food security and increasingly challenged by biotic stresses, particularly fungal diseases.

Among these challenges, rice blast disease, caused by the fungal pathogen *Magnaporthe oryzae* is considered one of the most destructive diseases of rice worldwide. This disease causes substantial yield losses of up to 30% with complete crop failure occurring under favourable environmental conditions (Asibi et al., 2019; Dean et al., 2012) by damaging several plant organs that impair the plant photosynthesis and grain production (Samal et al., 2020). with complete crop failure occurring under favourable environmental conditions. Climate change is expected to further increase the threat posed by rice blast. Rising temperatures, increased humidity, and altered rainfall patterns create favourable conditions for the development and spread of M. oryzae, particularly in the Himalayan foothills and adjoining Terai region (Wassmann et al., 2009; Singh and Maurya, 2021). These environmental changes may facilitate the emergence of new pathogen populations and increase disease incidence, posing additional challenges for rice production and food security (Cheng et al., 2020).

A major factor contributing to the persistence and widespread impact of rice blast is the remarkable genetic and phenotypic diversity of *M. oryzae*. Through mutation, recombination, and natural selection, the pathogen rapidly evolves, enabling it to adapt to changing environmental conditions and overcome resistance genes deployed in rice cultivars. This diversity is reflected in differences in morphology, growth characteristics, pathogenicity, and molecular profiles among isolates from different geographical regions. Consequently, isolates vary in their virulence and their ability to evade host defence mechanisms, with some capable of suppressing immune responses and causing severe disease, while others remain less aggressive or are restricted to specific cultivars (Ghatak et al., 2013; Chakraborty et al., 2020a; Dinkwar et al., 2023). Such isolate-specific interactions often result in the breakdown of host resistance, meaning that cultivars resistant in one region may become susceptible in another. Understanding the extent of local pathogen diversity and virulence is therefore essential for developing durable resistance through informed breeding strategies, including the deployment and pyramiding of effective resistance genes, as well as for designing region-specific disease management programmes (Panda et al., 2017; Gupta et al., 2018).

Regarding how *M. oryzae* spreads and evolves across landscapes, researchers employ risk distribution mapping and clustering analyses, which visualize disease severity, often quantified using indices like the Percent Disease Index (PDI), across different isolates (Khalaf et al., 2024). Clustering groups genetically or virulently similar isolates, often reflecting geographic origins, revealing potential migration routes and localized outbreak zones. These insights guide targeted surveillance and resource allocation, supporting strategies to mitigate epidemic spread (Rizzo et al., 2023; Eze et al., 2025). Accurate identification and evolutionary tracking of *M. oryzae* isolates, often using molecular markers like the Internal Transcribed Spacer (ITS) region of ribosomal DNA, allows construction of phylogenetic trees to uncover lineage diversification, gene flow, and virulence emergence (Gladieux et al., 2018; Patwardhan et al., 2014; Unemo and Shafer, 2014).

Despite the importance of rice blast in northern India, information on the morphological and molecular diversity of *M. oryzae* populations from the Terai foothills of the Himalayas remains limited. Therefore, this study aimed to characterize the morphological, cultural, pathogenic, and molecular diversity of *M. oryzae* isolates collected from this emerging agroecological region. Virulence screening was performed to identify highly aggressive isolates, followed by comparative analyses of colony morphology, conidial characteristics, growth behaviour on different culture media, and ITS-based phylogenetic relationships. The findings provide new insights into the diversity and evolution of *M. oryzae* populations in the Terai foothills and establish a foundation for region-specific resistance breeding and integrated disease management strategies.

## 2. Materials and Methods

### 2.1. Study area and sample collection

The *M. oryzae* isolates were collected from different rice fields across Terai Himalayan foothill areas known for their diverse rice cultivation systems during the 2023-2024 growing season. For this, rice plants exhibiting symptoms of rice blast disease, such as lesions on leaves, stems, and panicles (Supplementary Figure 1), were selected from fields in Assam, Bihar, and Uttar Pradesh, West Bengal including samples from both traditional Sali rice and floating rice varieties (Supplementary Table 1). These areas were chosen because of their high vulnerability to rice blast, making them ideal for pathogen isolation and characterization (Panda et al., 2017).

### 2.2. Pathogen isolation and maintenance

The isolation of *M. oryzae* from infected rice tissues was performed using standard methods (Gupta *et al*., 2020). Briefly, infected leaf tissues were excised into 5mm × 5mm sections and surface-sterilized using a 0.5% sodium hypochlorite solution for 2 minutes. After rinsing with sterile distilled water, the samples were placed on Potato Dextrose Agar (PDA) and incubated at 25°C in the dark for 5-7 days. Mycelium developing from the infected tissue segments was subcultured onto fresh PDA plates and incubated under the same conditions. The isolates were identified based on spore morphology (Supplementary Fig 2) and stored at 4°C until further use. The stock cultures were stored at 4°C for long-term preservation (Fajarningsih, 2016).

### 2.3. Screening of Isolates for disease severity

Isolates were inoculated on MTU-7029, a blast-susceptible rice variety, and grown under semi-controlled conditions (mean temperature: 28°C; relative humidity: >85%), simulating the Terai region environment in three independent trials (replicates). Disease severity was assessed using the Percent Disease Index (PDI) per isolate across three independent replicates, which were evaluated under identical conditions to ensure consistency and reproducibility of disease expression. Disease scoring was performed using the 0-9 IRRI-SES scale (IRRI, 2013) as detailed in Table 1. The PDI was calculated as follows:

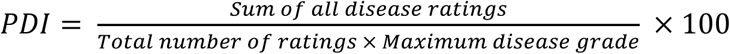

**Table 1.** Disease scoring scale IRRI-SES. Scale categories with their symptoms associated are represented

| Scale | Disease severity |
| --- | --- |
| 0 | Lesions are not present |
| 1 | Small brown specks of pinpoint size or larger brown specks without sporulation |
| 2 | Small roundish to slightly elongated, necrotic grey spots (1-2 mm in diameter) with a distinct brown margin. Lesions are mostly found on the lower leaves |
| 3 | Lesion type as in scale 2, but a significant number of lesions on upper leaf area |
| 4 | Typical susceptible blast lesions, >3 mm infecting less than 4% of leaf area |
| 5 | Typical susceptible blast lesions infecting 4-10% of the leaf area |
| 6 | Typical susceptible blast lesions infecting 11-25% of the leaf area |
| 7 | Typical susceptible blast lesions infecting 26-50% of the leaf area |
| 8 | Typical susceptible blast lesions infecting 51-75% of the leaf area and dead leaves |
| 9 | More than 75% leaf area affected |

The PDI values for each isolate across the three replicates were compiled and standardized using Z-score normalization to ensure consistency across replicates following:

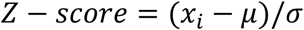

where, x_i_ = is the PDI value, μ= is the mean, and σ =is the standard deviation of the PDI values. Principal Component Analysis (PCA) was performed on the standardized PDI values to reduce dataset complexity and identify the most significant sources of variability (Wold *et al.,* 1987). K-means clustering was applied to the first two principal components to discriminate severity clusters. The optimal number of clusters (k) was determined using the elbow method based on within-cluster sum of squares. Based on consistently high PDI values across replicates, the two most aggressive isolates based on were selected for further detailed studies.

To visualise the spatial distribution of disease severity, geographic coordinates were assigned to each isolate based on its sampling location within the Terai belt and Himalayan foothill regions. A scatter map was generated in Python using Matplotlib, with longitude and latitude representing sampling locations. Marker size was scaled according to the average Percent Disease Index (PDI), allowing visualization of the spatial distribution of highly virulent isolates and potential disease hotspots having a putative risk distribution plotting.

### 2.4. Morphological characterization of severe isolates

#### Fungal growth

Six different media were used: ½ strength Potato Dextrose Agar (½ PDA), Potato Dextrose Agar (PDA), Oat Meal Agar (OMA), Rice Straw Extract Dextrose Oat Meal Agar (RSEDOMA), Rice Leaf Decoction (RLD) agar, and Mathur’s medium to evaluate the cultural characteristics and assess the growth patterns and adaptability of the pathogen on different media. Fungal cultures were incubated at 28°C for 10 days. Colony morphology was observed and recorded, including growth type, elevation, margin, colony colour, and surface texture.

Mycelial growth was measured by inoculating five 5-mm mycelial discs from each isolate into 250 ml sterilized conical flasks containing 100 ml Potato Dextrose Broth (PDB). Flasks were incubated at 28°C in a BOD incubator with shaking at 60 rpm. After 15 days, mycelia were harvested by filtering through pre-weighed Whatman filter paper No. 1. The filter paper and mycelium were dried at 50°C for 8 hours per day until constant weight was achieved. Dry mycelial weight was calculated by subtracting the initial filter paper weight.

The colony growth of *M. oryzae* was measured on at 5-, 10- and 15-days after inoculation (DAI) and the Area Under the Growth Progress Curve (AUGPC) was calculated to quantify the growth over time (Gupta et al., 2018; Ghatak et al., 2013) using the following formula:

Where:

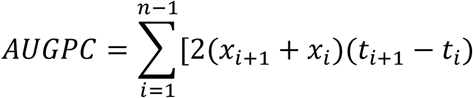

- X_i_ is the growth of the pathogen on the i^th^ date
- T_i_ is the time at which the i^th^ observation was recorded
- n is the number of days of observation

#### Spores germination and pathogenicity assays

Conidial germination was assessed at 6- and 24-hours post-inoculation. A 100 μL aliquot of conidial suspension (1 x 10⁵ conidia/mL) from each isolate was pipetted onto slides containing 2% water agar blocks. Slides were incubated at 28°C in Petri dishes lined with moist filter paper to maintain humidity. Germ tube lengths were measured using Moticam 2.0 software, and time-course photographs documented the germination process. Finally, length and width of spores were measured from infected lesions using Moticam 2.0 software attached to a binocular light microscope (20X objective, 10X eyepiece).

For the pathogenicity assay, stems of 20-day-old Oryza sativa plants were collected from the field and cut into 1 cm-long segments. Fifteen stem segments were placed in 50 mL Erlenmeyer flasks and sterilized at 1.4 kg cm⁻² for 1 h 30 min. Each flask was inoculated with two 5 mm-diameter mycelial plugs from 15-day-old cultures of each isolate and incubated at room temperature. Three stem segments were sampled at 10, 20, and 60 days after inoculation (DAI). Each segment was transferred to a test tube containing 1 mL of sterile distilled water, vortexed to dislodge the conidia, and the resulting suspension was collected. Conidial concentration was then determined using a haemocytometer.

### 2.5. Molecular characterisation of *M. oryzae* isolates

#### Redox enzymes profiling

Frozen leaf samples (100 mg) were ground using a Polytron homogenizer (Brinkmann Instruments) in an ice-cold 50 mM potassium phosphate buffer (pH 7.5) at a ratio of 1:20 (w/v). To inhibit protease activity, 1 mM phenylmethylsulphonyl fluoride (PMSF) was added. The resulting homogenates were centrifuged at 3500 rpm (approximately 2500 x g) for 10 minutes at 4 °C. The supernatants were then collected promptly and used immediately for enzyme activity assays, which were performed kinetically as outlined in the subsequent sections (Hermes-Lima and Storey, 1995).

Superoxide dismutase (SOD) activity (units per mg of protein) was measured by assessing the inhibition of nitro blue tetrazolium (NBT, 0.025 mM) reduction by superoxide radicals (O_2_) generated through the xanthine/xanthine oxidase system at 560 nm. The reaction mixture contained 50 mM sodium carbonate buffer, 0.1 mM xanthine, 0.025 mM NBT, 0.1 mM EDTA, and xanthine oxidase (0.1 U/ml), maintained at 25 °C. The reaction was initiated by adding a 1:20 (v/v) dilution of the tissue homogenate (Suzuki, 2000).

Catalase (CAT) activity (units per mg of protein) was assayed by monitoring the decrease in hydrogen peroxide (H_2_O_2_, 10 mM) concentration in 50 mM potassium phosphate buffer (pH 7.0) at 240 nm after adding the tissue extract. The assay components were maintained at 25 °C (Pippenger et al., 1998).

#### Phylogeny and evolution analysis

The selected highly virulent isolates (designated UBKV1 and UBKV2) were maintained on Potato Dextrose Agar (PDA) at 25°C for 7 days prior to DNA extraction. Genomic DNA was extracted using the DNeasy Plant Mini Kit (Qiagen, Hilden, Germany), following the manufacturer’s protocol. DNA quality and concentration were assessed using a Nanodrop spectrophotometer (Thermo Scientific, USA).

The internal transcribed spacer (ITS) region of ribosomal DNA was amplified using universal primers ITS1 and ITS4. Each 25 μl PCR reaction contained 1× PCR buffer, 2.0 mM MgCl₂, 0.2 mM dNTPs, 0.4 μM of each primer, 1 LA Taq DNA polymerase, and 50 ng of genomic DNA template. The PCR thermal cycling conditions were as follows: initial denaturation at 94°C for 5 min; 35 cycles of 94°C for 30 s, 55°C for 30 s, and 72°C for 45 s; followed by a final extension at 72°C for 7 min. The PCR products were purified using the QIAquick PCR Purification Kit (Qiagen, Germany) and sequenced bidirectionally using Sanger sequencing. Chromas Pro software (Technelysium Pty Ltd) was used to inspect sequence quality, and ambiguous regions were manually edited to ensure accuracy.

The obtained ITS sequences were curated and contigs were identified through BLAST searches NCBI against *Magnaporthe* genus, Later the programme Molecular Evolutionary Genetic Analysis Software, ver. 11.0 (MEGA 11.0) (K. Tamura et al., 2021) was used to edit and align the sequence files using CLUSTAL W algorithm (J.D Thompson et al., 1994). A phylogenetic tree was constructed using the maximum likelihood method in MEGA 11.0, incorporating 31 isolates, including annotations for *M. grisea* and one outgroup (*Fusarium oxysporum*, accession no. MZ496570). Confidence levels were assessed using 1000 bootstrap replicates.

Principal Component Analysis (PCA) and Multidimensional Scaling (MDS) were performed on the standardized pairwise similarity matrix using Python 3.10 with scikit-learn and matplotlib libraries (Pedregosa et al., 2011). MDS was applied to the genetic distance matrix using the MDS function in scikit-learn, Python (scikit-learn, matplotlib) v1.2.2 with Euclidean distance metric, two dimensions, random initialization, and a maximum of 300 iterations. Silhouette analysis was performed using the silhouette score function in scikit-learn with Euclidean distance to evaluate cluster quality. The criteria followed was: Bootstrap confidence ≥70% for robust phylogenetic support, Silhouette score ≥0.25 indicating moderate clustering, MDS stress value <0.1 indicating a good fit, PCA explained variance ≥60% considered acceptable (observed 68.9%), SPSS and R studio used for Data analysis and visualization.

## 3. Results

### 3.1 Pathogenicity assessment and spatial distribution of *Magnaporthe oryzae* isolates

A total of 48 M. oryzae isolates were recovered from rice leaf samples collected across the Terai foothills of the Himalayas in Assam, Bihar, Uttar Pradesh, and West Bengal. All isolates grew consistently on PDA, producing colonies that were initially white to greyish and later developed characteristic concentric dark pigmentation. Microscopic examination confirmed typical M. oryzae morphology, including septate hyphae and pyriform, two-septate conidia. Pathogenicity screening revealed substantial variation in disease severity among the isolates, with Percent Disease Index (PDI) values ranging from 22.2% to 64.1% (Fig. 1). The least virulent isolates originated from Buxar (29.7 ± 2.8%) and Kalimpong (28.7 ± 3.5%), whereas isolates from Alipurduar (64.9 ± 2.3%) and Siliguri (63.8 ± 2.5%) exhibited the highest disease severity.

**Figure 1.**
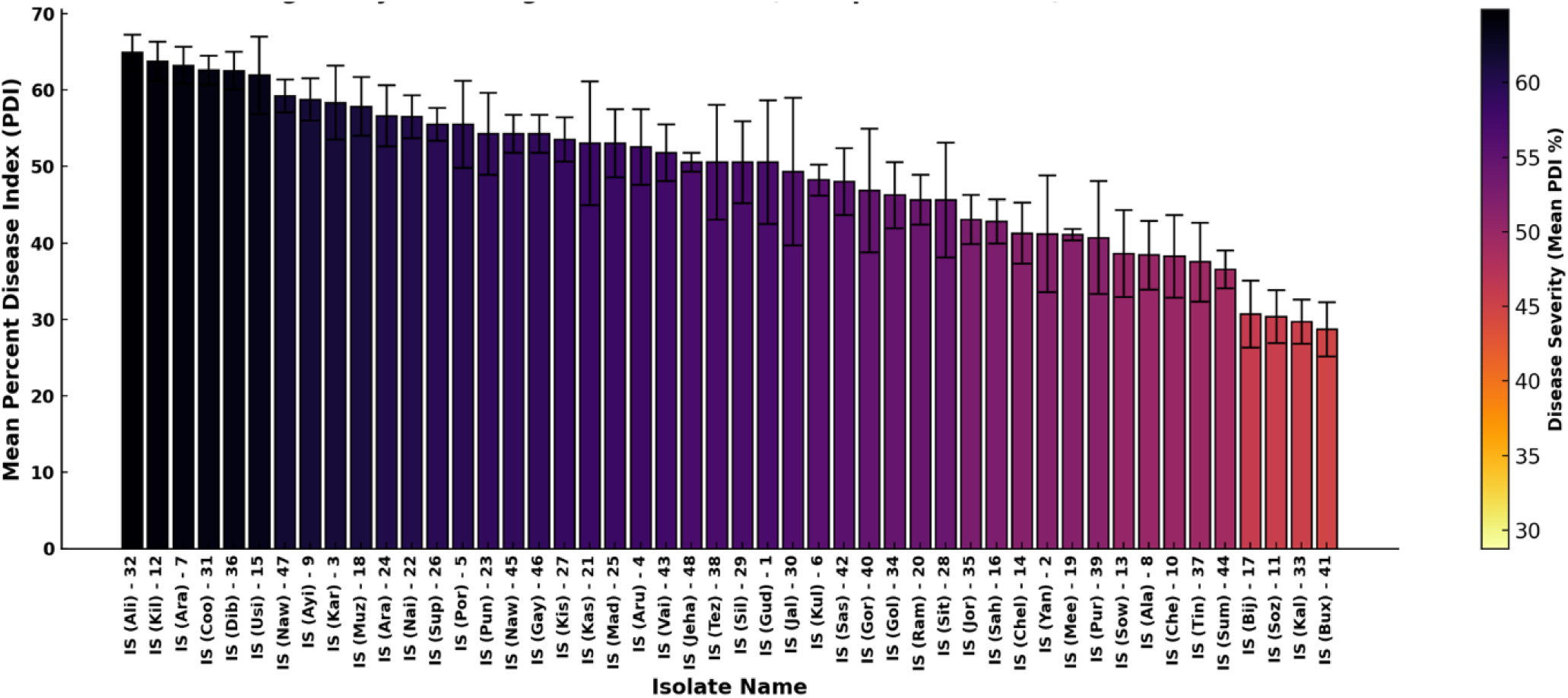
Pathogenicity screening of 48 *M. oryzae* isolates. Mean Percent Disease Index (PDI ± SE) of isolates ranked from highest to lowest virulence. Disease severity groups were determined by one-way ANOVA followed by Tukey’s HSD test.

Principal component analysis (PCA) of standardized PDI values explained 76.4% of the total variation, with PC1 and PC2 accounting for 47.5% and 28.9%, respectively (Fig. 2A). K-means clustering (k = 3) classified the isolates into three pathogenicity groups that closely corresponded to their PDI rankings. Cluster 1 comprised highly pathogenic isolates (PDI > 60%), including IS (Ali)-32, IS (Kil)-12, IS (Ara)-7, IS (Coo)-31, and IS (Dib)-36, and exhibited the greatest within-cluster variability, indicating considerable phenotypic diversity among highly virulent isolates. Cluster 2 included moderately pathogenic isolates (PDI 45–60%), such as IS (Muz)-18, IS (Naw)-47, IS (Sup)-26, and IS (Aru)-4, with intermediate levels of disease severity and variability. In contrast, Cluster 3 consisted of weakly pathogenic isolates (PDI < 35%), represented by IS (Bux)-41, IS (Kal)-33, IS (Soz)-11, and IS (Bij)-17, and formed a tightly grouped cluster, reflecting low variability and consistently reduced pathogenicity. Ward’s hierarchical clustering produced a comparable grouping pattern with high intra-cluster similarity and clear separation among clusters, confirming the robustness of the pathogenicity classification. The hierarchical plot also revealed that the intra-cluster similarity was high, and inter-cluster distances were large, further confirming that the three identified groups are biologically meaningful.

**Figure 2.**
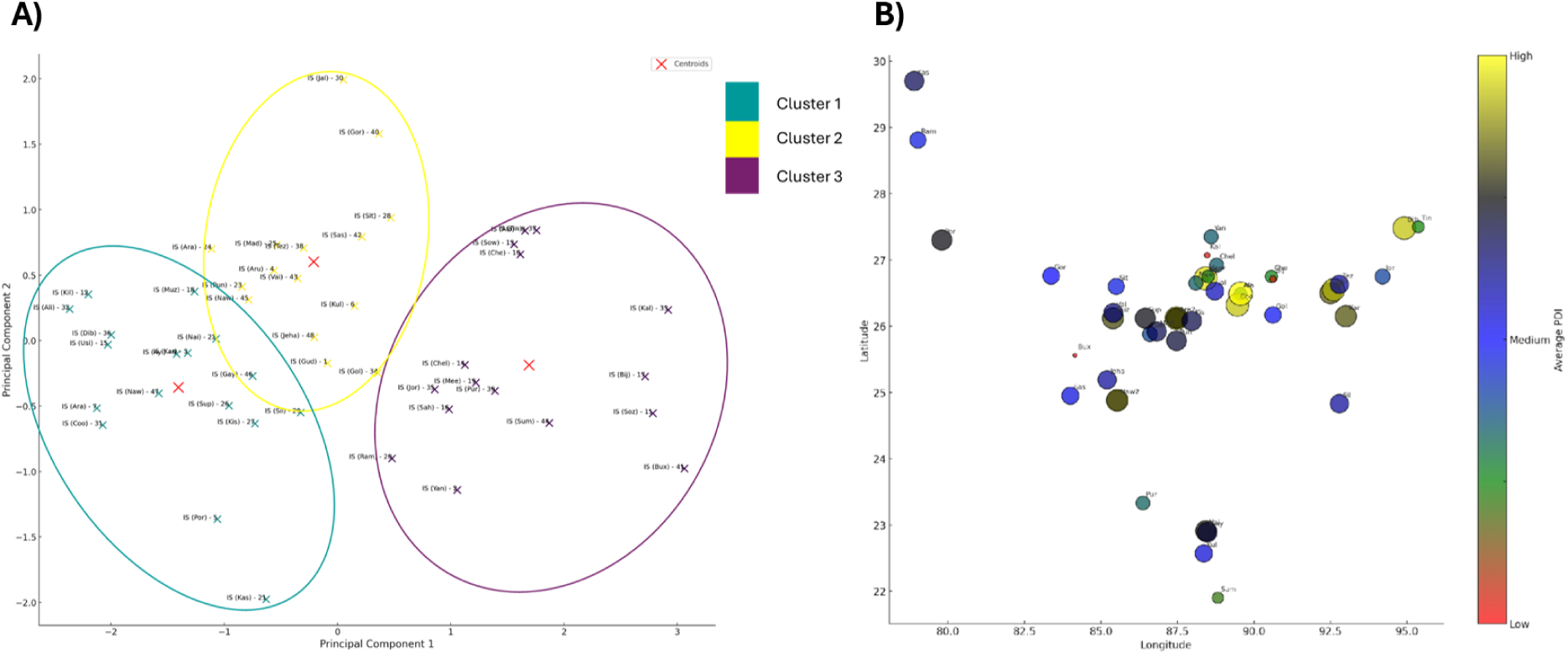
Principal Component Analysis (PCA) and Risk Zone Prediction of 48 *M. oryzae* Isolates. (**A**) PCA plot showing the clustering of isolates based on pathogenic and morphological traits using K-means clustering (k = 3); red crosses indicate cluster centroids. (**B**) Spatial distribution of isolates according to sampling locations, with bubble size representing disease intensity and colour indicating the average percent disease index (PDI), illustrating regional disease risk.

Spatial mapping of PDI values identified disease severity hotspots within the Terai belt and Himalayan foothill regions (Fig. 2B). The highest disease severity was recorded for isolates collected from Alipurduar (64.9%), Siliguri (63.8%), Cooch Behar (≈63%), Araria (≈63%), and Dibrugarh (≈62%), whereas neighbouring districts generally exhibited moderate disease severity (40–60%). The geographic distribution of highly virulent isolates closely corresponded to Cluster 1 identified by the PCA and K-means analyses, and Pearson correlation analysis showed a strong positive association between cluster membership and disease severity (r = 0.78). Although based on observed pathogenicity rather than predictive modelling, the spatial distribution identifies areas where highly virulent M. oryzae populations are concentrated and therefore highlights regions with an increased potential for disease outbreaks under favourable environmental conditions. These findings provide a valuable framework for prioritizing disease surveillance, targeted monitoring, and region-specific deployment of resistant rice cultivars across the Terai foothills and adjoining Himalayan regions.

### 3.2. Differential growth and sporulation characteristics of UBKV1 and UBKV2

To further characterise the two most virulent isolates, UBKV1 (Siliguri) and UBKV2 (Alipurduar), detailed morphological and molecular analyses were performed. Conidial morphology (Fig. 3) revealed significant differences between the isolates. UBKV1 produced conidia with a mean length of 28.07 µm and a width of 12.57 µm, whereas UBKV2 produced significantly longer but narrower conidia, with a mean length of 33.61 µm and a width of 10.59 µm (Fig. 3A, B). The difference in conidial length was highly significant (p = 4.16 × 10⁻⁵), suggesting morphological divergence between the two highly virulent isolates. In addition, UBKV2 exhibited significantly greater germ tube elongation, with a mean germ tube length of 50.41 µm compared with 43.86 µm for UBKV1 (p = 5.2 × 10⁻⁴) (Fig. 3C).

**Figure 3.**
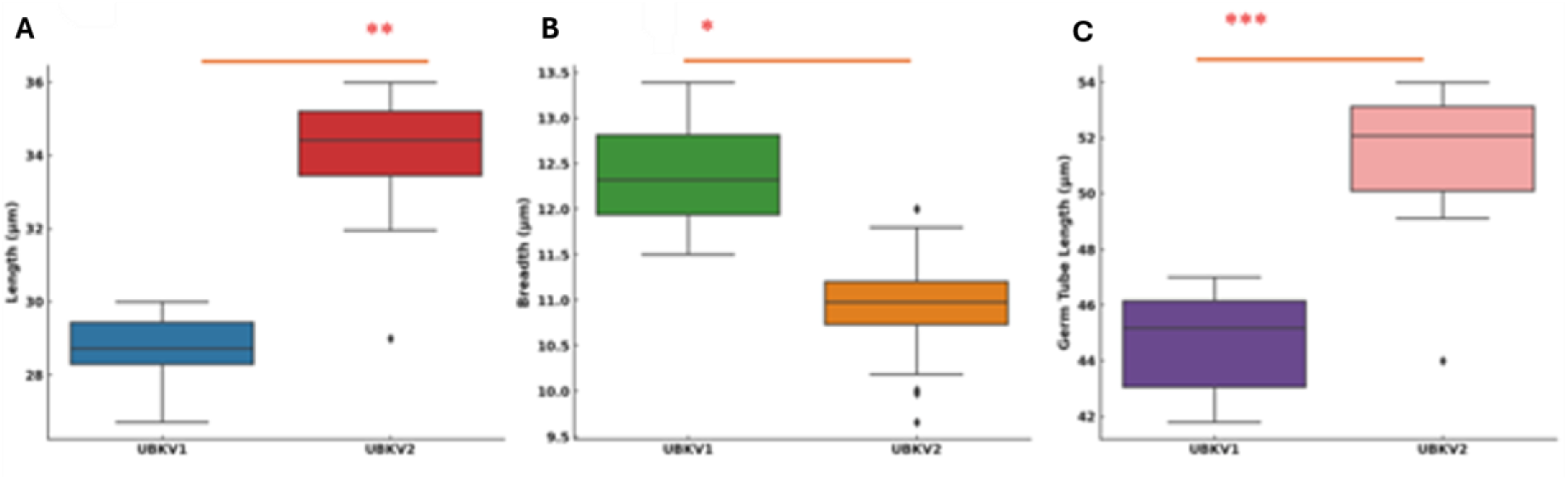
Comparative morphological analysis of conidial structures between the most virulent isolates (UBKV1 and UBKV2). (**A**) Boxplot showing the comparison of conidial length (µm) between isolates UBKV1 and UBKV2. (**B**) Boxplot representing the conidial breadth (µm) comparison between the two isolates, showing a significant difference (p < 0.05, denoted by *). (**C**) Boxplot depicting the germ tube length (µm) of UBKV1 and UBKV2 (p < 0.001, denoted by ***). All morphological measurements were based on microscopic evaluations of conidia grown under identical laboratory conditions. Statistical analysis was performed using Student’s t-test.

Fresh mycelial biomass differed significantly between the two isolates (Fig. 4A). UBKV2 produced a significantly greater fresh mycelial weight (22.98 g) than UBKV1 (15.82 g), indicating a higher vegetative growth rate under identical culture conditions. In contrast, dry mycelial weight showed only a slight, non-significant difference (p = 0.157), with UBKV2 and UBKV1 producing 1.31 and 1.20 g, respectively (Fig. 4B). These results suggest that although UBKV2 accumulated greater fresh biomass, both isolates produced comparable amounts of dry fungal biomass.

**Figure 4.**
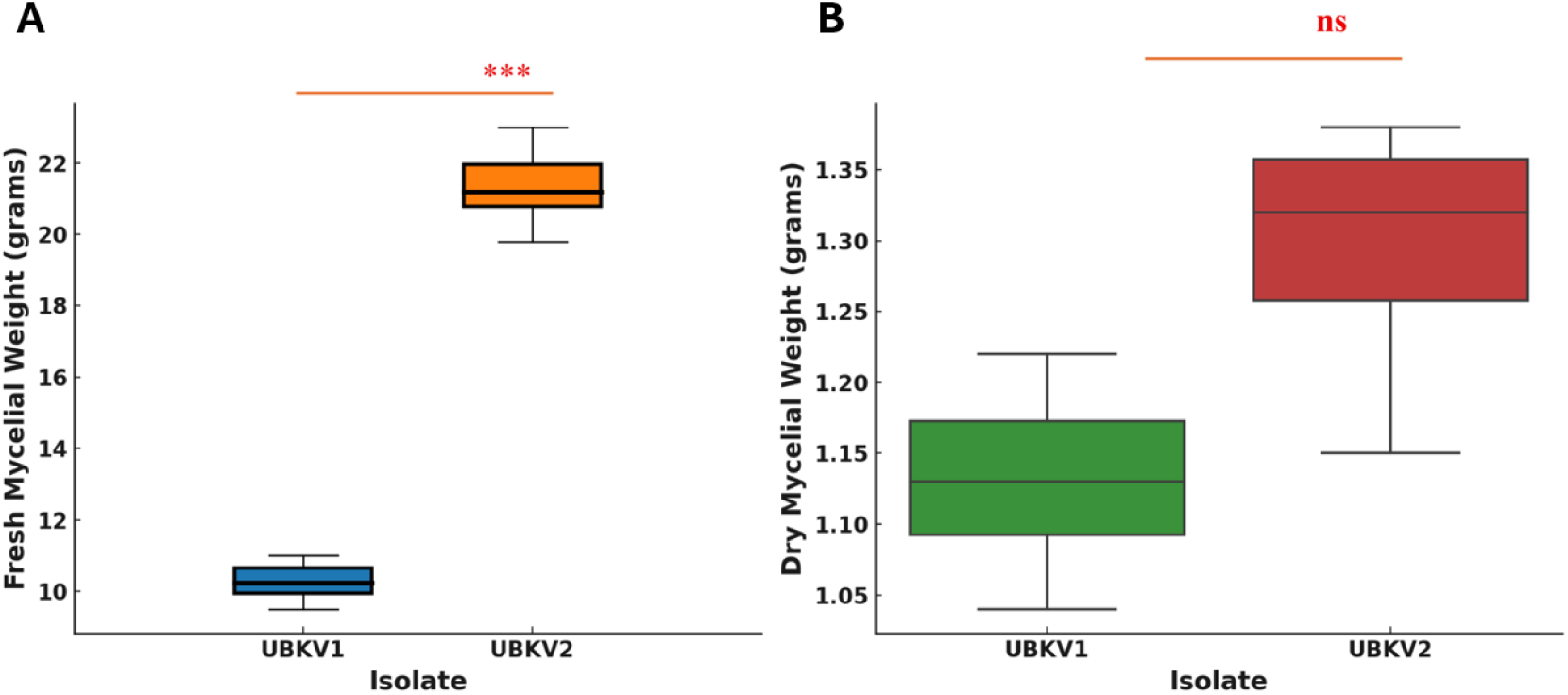
Comparison of mycelial biomass between Isolates UBKV1 and UBKV2. Boxplots showing (**A**) fresh and (**B**) dry mycelial weight. Fresh biomass differed significantly between isolates (***, *p* < 0.001), whereas dry biomass showed no significant difference (ns). Statistical significance was determined using Student’s *t*-test.

Cultural characteristics on six growth media (Fig. 5) further distinguished the two isolates. On PDA and OMA, both isolates produced typical cottony, white colonies; however, UBKV1 formed slightly larger colonies with more abundant aerial mycelium, particularly on RLD medium. Colony pigmentation varied with the growth medium, changing from white to greyish or light brown on PDA and OMA, whereas darker central pigmentation developed on Mathur’s medium and RSEDOMA. Colony morphology also differed among media: both isolates formed regular colonies with dull white to milky white mycelia and indistinct zonation on PDA and OMA, while growth was more restricted on Mathur’s medium and RSEDOMA. Under these conditions, UBKV1 exhibited more pronounced sectoring and darker pigmentation than UBKV2. On RLD medium, both isolates developed regular colonies with a greyish centre that darkened with incubation. Moreover, radial growth measurements (Fig. 5) showed that UBKV1 consistently grew faster than UBKV2 across all media tested, with the greatest colony diameters recorded on PDA and Mathur’s medium. After 7 days, colony diameters reached 6.5 cm for UBKV1 and 5.8 cm for UBKV2 on PDA. These differences were significant according to the LSD test (LSD = 0.0146). In contrast, both isolates exhibited the slowest growth on RLD medium, demonstrating the influence of culture medium on fungal growth.

**Figure 5.**
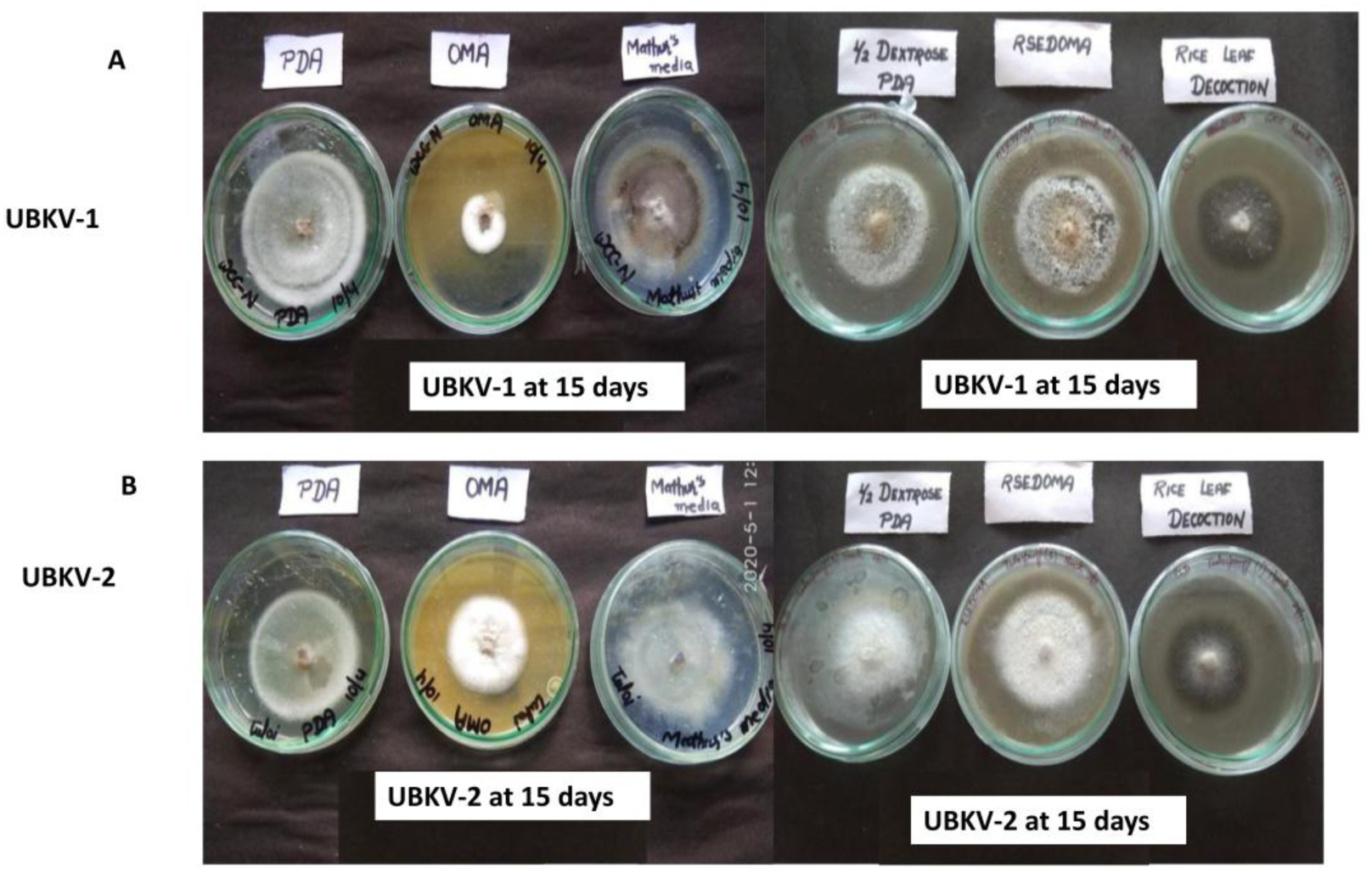
Growth rate comparison and cultures characteristics of *M. oryzae* isolates (UBKV1 and UBKV2) grown on different media after 15 days of incubation across six different culture media. (**A**) Colony morphology of isolate UBKV-1 on Potato Dextrose Agar (PDA), Oatmeal Agar (OMA), Mathur’s medium, ½ Dextrose PDA, RSEDOMA, and Rice Leaf Decoction media after 15 days of incubation at 28 ± 2 °C. (**B**) Colony morphology of isolate UBKV-2 grown under the same conditions and media for comparison.

Sporulation and conidial production studies showed that both isolates produced abundant conidia on PDA, OMA, and RLD after 7 days of incubation. The conidia were cylindrical, smooth, and transparent, consistent with typical *M. oryzae* morphology. Interestingly, UBKV1 exhibited slightly higher conidial density, averaging 2.5 × 10⁶ spores/mL, compared to 2.2 × 10⁶ spores/mL for UBKV2. The Area Under the Growth Progress Curve (AUGPC) (Fig.6) was used to integrate temporal growth dynamics across the media. Results indicated that UBKV1 consistently exhibited higher AUGPC values than UBKV2 on all tested media, with the highest values recorded on Mathur’s medium and PDA. UBKV2’s highest AUGPC was observed on ½ Strength PDA, suggesting that it might be better adapted to suboptimal nutrient conditions. Statistically significant differences in AUGPC values confirmed that UBKV1 outperformed UBKV2 in terms of cumulative growth and pathogenic potential under the tested conditions.

**Figure 6.**
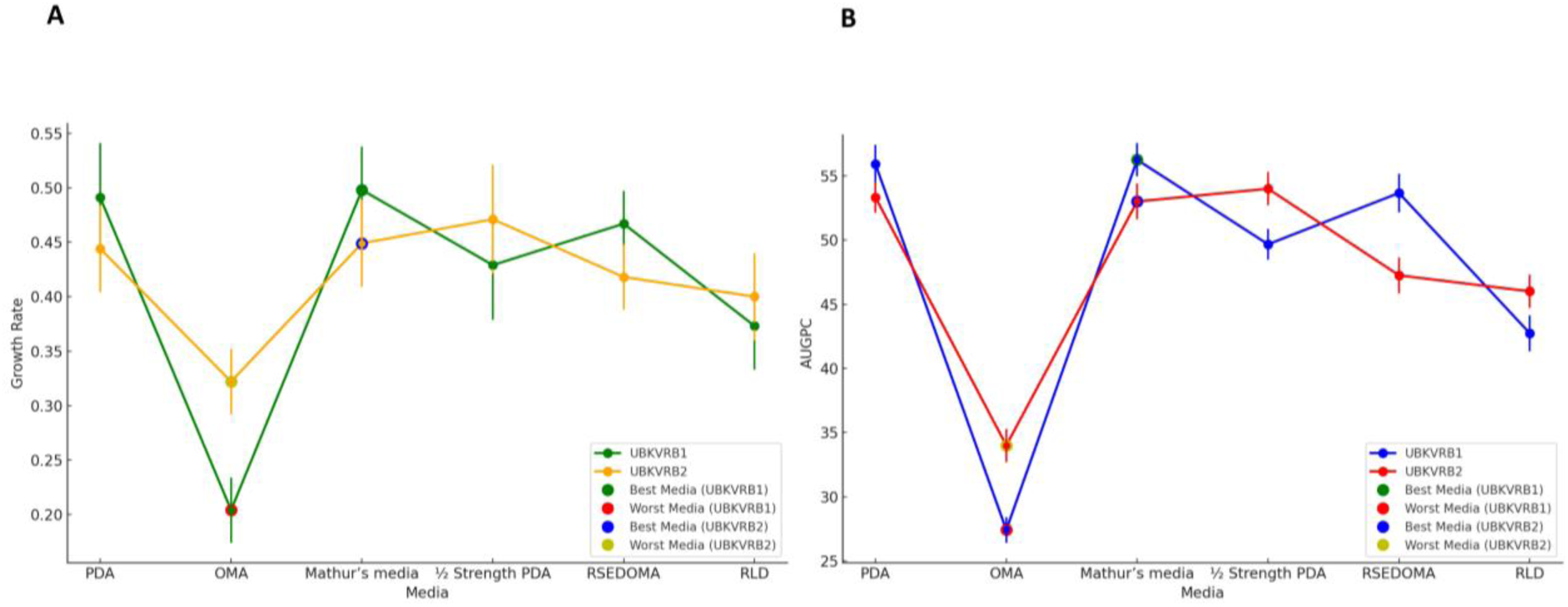
Growth performance of isolates UBKVRB1 and UBKVRB2 on different culture media. (**A**) Line plot showing the growth rate (cm/day) of isolates UBKVRB1 and UBKVRB2 cultured on six different media (PDA, OMA, Mathur’s medium, ½ Strength PDA, RSE DOMA, and RLD). The best- and worst-performing media for each isolate are marked with coloured indicators. (**B**) Line plot illustrating the Area Under Growth Curve (AUGC) values of the same isolates across the same media.

On water agar, both *M. oryzae* isolates produced germ tubes that underwent developmental switches (ds) to form preliminary invasive hyphae (Pre-IH), although appressoria did not develop due to the absence of host-derived cues. Germ tube elongation increased progressively from 0 to 24 h, with UBKV2 consistently producing longer germ tubes than UBKV1. Conidia of UBKV1 germinated from one, two, or all three cells, whereas UBKV2 germinated from one or two cells only. Developmental switches leading to Pre-IH were observed after 24 h in both isolates, with a slightly earlier onset in UBKV2 (Fig. 7).

**Figure 7.**
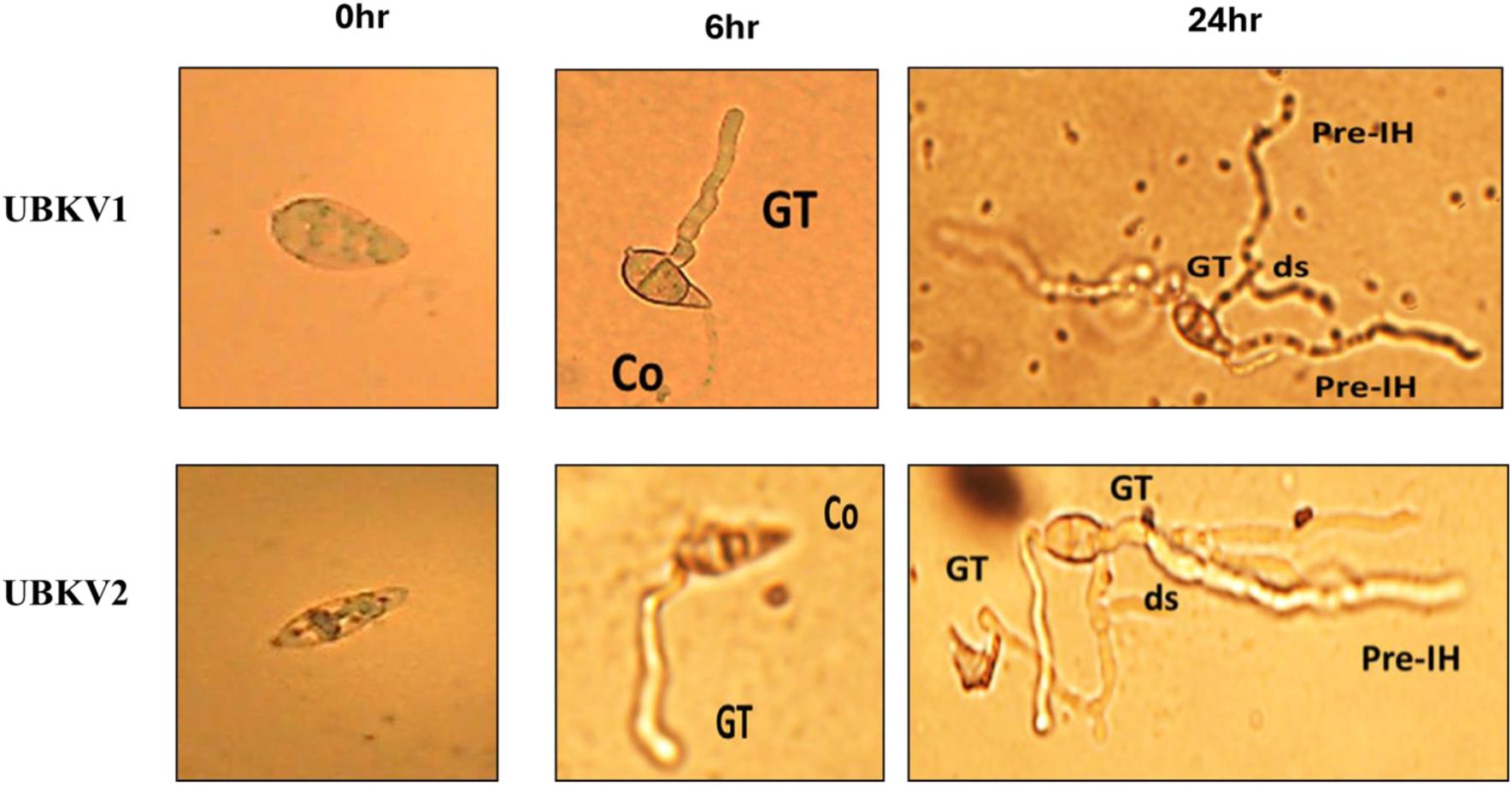
Conidial germination and pre-infection structure development in *M. oryzae* isolates UBKV1 and UBKV2. Representative light micrographs showing conidial germination after 0, 6, and 24 h of incubation on water agar. (**A–C**) UBKV1: (**A**) resting conidium (Co); (**B**) germ tube (GT) emergence; (**C**) preliminary invasive hyphae (Pre-IH) with developmental switch (ds). (**D–F**) UBKV2: (**D**) resting conidium (Co); (**E**) germ tube (GT) elongation; (**F**) Pre-IH formation with developmental switch (ds). Images were captured under bright-field microscopy (40×) after incubation at 25 ± 1 °C. Abbreviations: Co, conidium; GT, germ tube; ds, developmental switch; Pre-IH, preliminary invasive hyphae.

Spore density increased progressively in both isolates over time, although their sporulation patterns differed (Fig. 8). At 10 DAI, both isolates exhibited low spore densities (<1,000 spores mL⁻¹). Sporulation increased substantially by 20 DAI, with UBKV1 producing higher spore densities than UBKV2. This difference became more pronounced at 60 DAI, when UBKV1 exhibited the highest sporulation, whereas UBKV2 maintained comparatively lower spore densities. UBKV1 also showed greater variability in sporulation than UBKV2.

**Figure 8.**
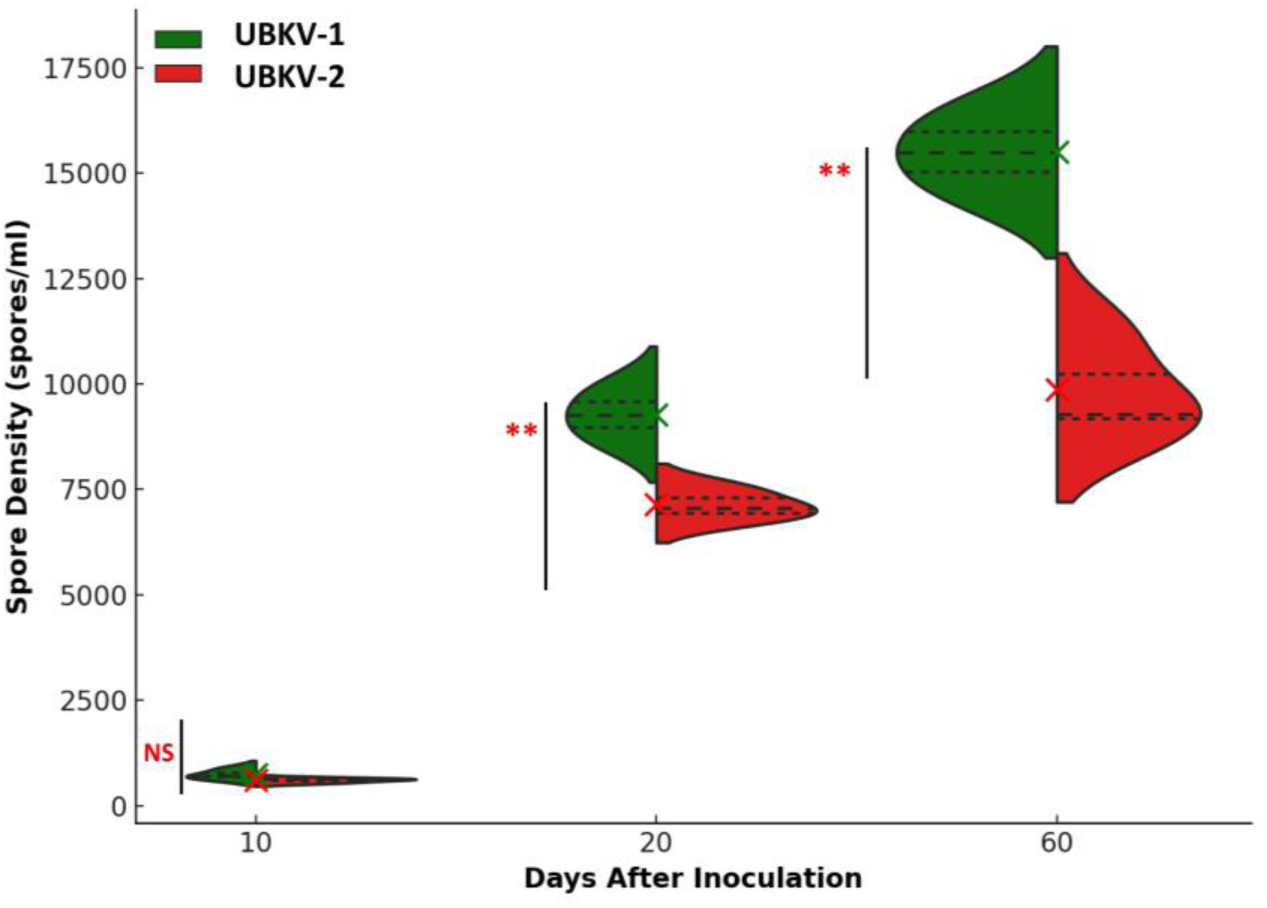
Temporal spore density distribution of Isolates UBKVRB1 and UBKVRB2. Violin plots showing spore density (spores mL⁻¹) at 10, 20, and 60 DAI. Violin shapes represent data distribution, dotted lines indicate quartiles, and × denotes the mean. Statistical differences were analysed by one-way ANOVA followed by Tukey’s HSD test.

### 3.3. Redox Activity in Rice Under UBKV isolates Infection

Superoxide dismutase (SOD) and catalase (CAT) activities were assessed in healthy plants and plants inoculated with UBKV1 or UBKV2 at 24, 48, and 96 h post-inoculation (Fig. 9). Both isolates induced significantly higher SOD activity than the healthy control at 24 h (UBKV1: 45 U mg⁻¹; UBKV2: 40 U mg⁻¹; control: 26 U mg⁻¹) and 48 h (38 and 42 U mg⁻¹ vs. 30 U mg⁻¹; p < 0.05). By 96 h, SOD activity had declined to levels comparable with the control, although the decline was more rapid in UBKV1 than in UBKV2. Similarly, CAT activity was markedly elevated in both infected plants at 24 h (15.0 and 14.5 U mg⁻¹ vs. 3.5 U mg⁻¹ in the control; p < 0.001) and remained high at 48 h, with UBKV2 exhibiting significantly greater activity than UBKV1 (19 vs. 16 U mg⁻¹; p = 0.022). At 96 h, CAT activity decreased in both isolates but remained significantly higher in UBKV2 (14.5 U mg⁻¹) than in UBKV1 (7.0 U mg⁻¹) and the control (6.5 U mg⁻¹), indicating a more sustained antioxidant response in plants infected with UBKV2.

**Figure 9.**
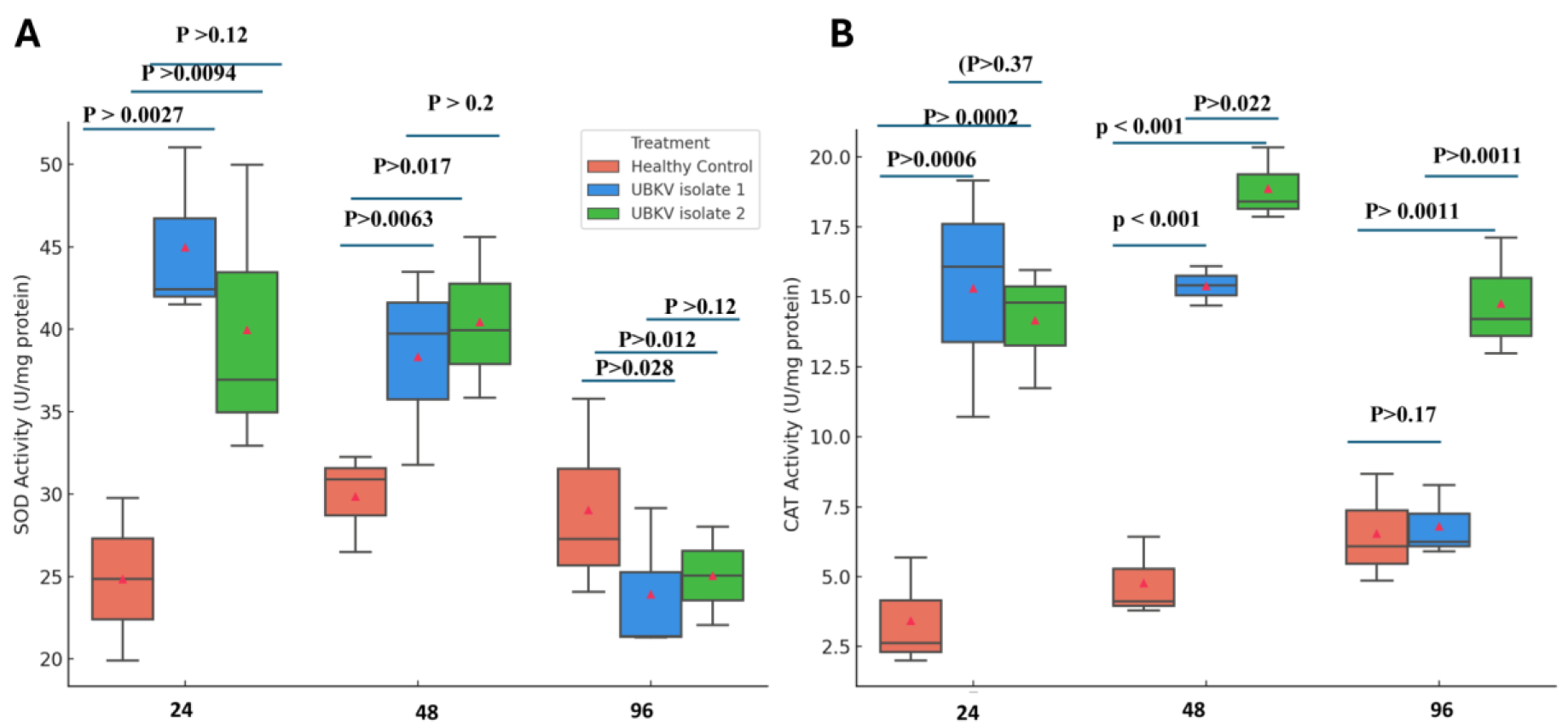
Antioxidant enzyme activities in rice plants inoculated with *M. oryzae* isolates UBKV1 and UBKV2. Boxplots showing (**A**) superoxide dismutase (SOD) and (**B**) catalase (CAT) activities at 24, 48, and 96 h after inoculation, compared with healthy controls. Red triangles (▴) indicate mean values. Statistical differences were determined by two-way ANOVA followed by Tukey’s HSD test (p < 0.05).

### 3.4. Phylogenetic analysis reveals distinct evolutionary lineages of UBKV1 and UBKV2

To investigate the genetic relationships of the two highly virulent isolates, ITS sequences from UBKV1 (MW882140) and UBKV2 (MW882141) were analysed together with 31 reference isolates of the *Magnaporthe oryzae/Pyricularia oryzae* complex representing diverse geographical origins across India and Asia, using *Fusarium oxysporum* (MZ496570) as the outgroup. Pairwise genetic distances among *M. oryzae* isolates ranged from 0.006 to 0.040, whereas the outgroup showed substantially greater divergence (0.28-0.31), confirming its suitability for rooting the phylogenetic tree.

Maximum Likelihood phylogenetic analysis using the Tamura-Nei substitution model with 1,000 bootstrap replicates recovered four major clades within the *M. oryzae* population, with bootstrap support values ranging from 46% to 99% (Fig. 10A). UBKV2 clustered within the eastern Indian lineage together with isolates from Odisha, exhibiting relatively short branch lengths consistent with high sequence similarity. In contrast, UBKV1 formed a distinct branch associated with isolates from southern India and Southeast Asia, displaying greater genetic divergence from the eastern Indian lineage.

**Figure 10.**
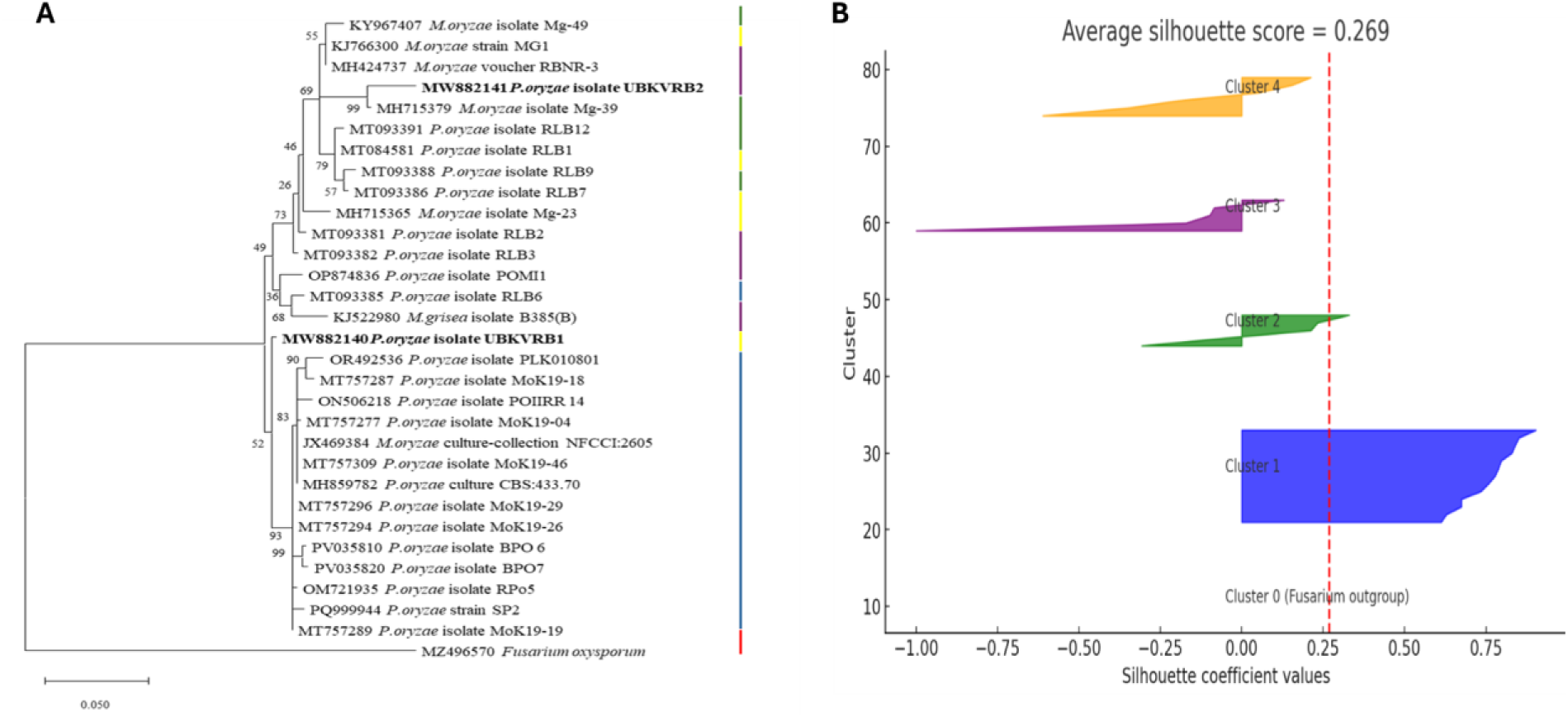
Figure 10. Phylogenetic relationships and cluster validation of *M. oryzae* isolates. (**A**) Maximum Likelihood phylogenetic tree based on ITS sequences (Tamura-Nei model; 1,000 bootstrap replicates), rooted with *Fusarium oxysporum*. (**B**) Silhouette analysis showing the clustering quality of individual isolates based on ITS sequence similarity.

To further evaluate genetic relationships, principal component analysis (PCA), multidimensional scaling (MDS), and k-means clustering were performed using ITS sequence similarity data. Five genetic clusters were identified, clearly separating *F. oxysporum* from the *M. oryzae* complex while revealing limited but detectable genetic variation among *M. oryzae* isolates. Cluster validation by silhouette analysis (Fig. 10B) yielded a mean coefficient of 0.269, indicating moderate separation among clusters. The outgroup formed a completely distinct cluster (silhouette = 1.00), whereas UBKV1 (0.40) and UBKV2 (0.35) were assigned to separate, well-defined clusters despite their close genetic relationship. Together, the phylogenetic, genetic distance, and clustering analyses consistently demonstrate that UBKV1 and UBKV2 represent genetically differentiated evolutionary lineages within the *M. oryzae* population.

## 4. Discussion

Rice blast remains one of the greatest constraints to sustainable rice production because *Magnaporthe oryzae* populations continuously evolve in response to host resistance and environmental selection. The present study provides the first integrated characterization of *M. oryzae* populations from the Terai Himalayan foothills by combining pathogenicity screening, morpho-cultural characterization, antioxidant responses, and molecular phylogenetics. Together, these complementary approaches revealed substantial phenotypic and genetic diversity among isolates and identified highly virulent populations concentrated within the Terai belt, emphasizing the importance of regional surveillance for disease forecasting and resistance breeding.

The resulting pathogenicity assays demonstrated considerable variation among the 48 isolates, with PDI values ranging from 22.2 to 64.9%, confirming the heterogeneous nature of M. *oryzae* populations reported in other rice-growing regions (Dean et al., 2012; Gladieux et al., 2018). PCA and clustering analyses separated the isolates into three well-defined pathogenicity groups that closely reflected disease severity, while spatial mapping identified Alipurduar, Siliguri, Cooch Behar, Araria, and Dibrugarh as major disease hotspots. The strong association between cluster membership and disease severity (r = 0.78) suggests that combining multivariate analyses with geographic information provides a practical framework for identifying regions at greater epidemic risk. Because the Terai belt is characterized by prolonged humidity, moderate temperatures, and extensive movement of rice germplasm, these highly virulent populations may act as important inoculum reservoirs capable of facilitating regional disease outbreaks under favourable environmental conditions (Savary et al., 2019; Bebber et al., 2014). Such information is particularly valuable for prioritizing surveillance efforts and deploying resistant cultivars according to local pathogen populations.

Detailed characterization of the two most aggressive isolates revealed contrasting biological strategies associated with virulence. UBKV2 (Alipurduar) produced significantly longer conidia (33.61 µm vs. 28.07 µm), longer germ tubes (50.41 vs. 43.86 µm), and greater fresh mycelial biomass (22.98 vs. 15.82 g) than UBKV1, indicating greater vegetative growth and potentially faster host colonization. In contrast, UBKV1 exhibited faster radial growth across all culture media, higher cumulative growth (AUGPC), and consistently greater sporulation, producing approximately 2.5 × 10⁶ spores mL⁻¹ compared with 2.2 × 10⁶ spores mL⁻¹ for UBKV2 and maintaining higher spore densities throughout the incubation period. These contrasting traits suggest distinct fitness strategies, whereby UBKV2 may invest preferentially in vegetative growth and infection establishment, whereas UBKV1 maximizes inoculum production and dispersal. Similar trade-offs between colony growth, sporulation efficiency, and pathogenic fitness have been reported in geographically distinct populations of M. oryzae (Talbot, 2003; Wilson and Talbot, 2009), highlighting the remarkable phenotypic plasticity that underpins the adaptive success of this pathogen.

The antioxidant enzyme analyses further demonstrated that rice rapidly activates oxidative defence responses following infection, with marked increases in both SOD and CAT activities during the early stages of host-pathogen interaction. SOD activity increased from 26 U mg⁻¹ in healthy plants to 45 and 40 U mg⁻¹ following inoculation with UBKV1 and UBKV2, respectively, at 24 h post-inoculation, before gradually declining to near-control levels by 96 h. A similar transient pattern was observed for CAT, although UBKV2 induced a significantly stronger and more sustained response, maintaining CAT activity at 14.5 U mg⁻¹ after 96 h compared with only 7 U mg⁻¹ in UBKV1. The oxidative burst represents one of the earliest components of plant innate immunity, serving both as a direct antimicrobial mechanism and as a signalling pathway for defence activation (Torres et al., 2006; Apel and Hirt, 2004). However, successful pathogens have evolved sophisticated antioxidant systems and secreted effectors that detoxify reactive oxygen species or suppress ROS-mediated signalling, thereby facilitating host colonization (Egan et al., 2007; Wang et al., 2019). The sustained CAT response induced by UBKV2 therefore suggests a more prolonged oxidative interaction between host and pathogen and may reflect enhanced ROS detoxification during infection. These observations are consistent with studies showing that catalase activity and ROS homeostasis are critical determinants of M. oryzae pathogenicity and successful biotrophic growth (Skamnioti and Gurr, 2007; Ryder et al., 2013).

Our phylogenetic analyses supported the phenotypic observations by demonstrating that the two highly virulent isolates belong to distinct evolutionary lineages despite sharing a common species background. PCA, silhouette analysis, and Maximum Likelihood phylogeny consistently separated UBKV1 and UBKV2 into different genetic clusters. UBKV2 grouped within the eastern Indian lineage together with Odisha isolates and exhibited relatively short branch lengths, suggesting recent diversification within this regional population. In contrast, UBKV1 clustered with isolates from southern India and Southeast Asia and displayed greater genetic divergence, indicating an independent evolutionary trajectory. These findings agree with previous population genomic studies demonstrating that *M. oryzae* evolves through repeated cycles of mutation, migration, recombination, and local adaptation, generating geographically structured populations with distinct virulence profiles (Gladieux et al., 2018; Zhong et al., 2018). Although ITS sequences provide limited phylogenetic resolution compared with genome-wide analyses, the concordance between molecular, morphological, and pathogenic data supports the existence of biologically meaningful differentiation among the Terai isolates.

Collectively, these findings demonstrate that rice blast populations in the Terai Himalayan foothills comprise genetically and phenotypically diverse pathogen populations that differ not only in pathogenicity but also in growth strategy, host interaction, and evolutionary history. Integrating pathogenicity phenotyping, spatial epidemiology, morpho-cultural characterization, antioxidant profiling, and molecular phylogenetics provides a comprehensive framework for identifying emerging high-risk populations and supports the development of region-specific disease surveillance, resistance breeding, and integrated management strategies. Future studies incorporating whole-genome sequencing, effector profiling, and long-term epidemiological monitoring will further improve understanding of pathogen evolution and strengthen predictive approaches for sustainable rice blast management under changing climatic conditions.

## 5. Conclusion

This study provides the first integrated assessment of the morphological, pathogenic, molecular, and spatial diversity of *Magnaporthe oryzae* populations in the Terai Himalayan foothills of India. The identification of distinct virulence groups, geographically defined disease hotspots, and two highly aggressive isolates belonging to separate evolutionary lineages demonstrates that pathogen diversity is structured at both phenotypic and genetic levels within this emerging rice-growing region. These findings improve our understanding of blast epidemiology in the eastern Himalayan foothills and provide practical guidance for regional resistance breeding, targeted surveillance, and deployment of location-specific disease management strategies. As climate change is expected to accelerate pathogen evolution and disease spread, continuous monitoring of population structure and virulence will be critical for developing durable blast resistance and ensuring sustainable rice production in this strategically important agroecosystem.

## Conflict of interest

The authors declare that they have no known competing financial interests or personal relationships that could have appeared to influence the work reported in this paper.

## Authors’ contribution

Conceptualization of research (SJ, SB, AKC, KS); Designing of the experiments and contribution of experimental materials (SJ, SB); Execution of lab experiments (SJ, YMB, DS, RM); Analysis of data and interpretation (SJ, YMB, DS, RSL); Preparation and writing of the manuscript draft (SJ, YMB); Manuscript writing, review and editing (YMB and RSL); Final approval and submission (RSL).

## Acknowledgement

Authors gratefully acknowledge Uttar Banga Krishi Viswavidyalaya, Cooch Behar, West Bengal, India, for providing the providing all the necessary facilities to carry out the study.

## Notes

### Competing Interest Statement

The authors have declared no competing interest.

